# The adaptation of chlamydiae to facultative host multicellularity

**DOI:** 10.1101/2024.11.28.625222

**Authors:** Lukas Helmlinger, Patrick Arthofer, Norbert Cyran, Astrid Collingro, Matthias Horn

## Abstract

The phylum Chlamydiota consists of obligate intracellular bacteria comprising the human pathogen *Chlamydia trachomatis* and a large variety of species infecting animals and protists. Despite their enormous diversity, a feature shared by all known chlamydiae is their biphasic developmental cycle, consisting of intra- and extracellular stages with substantial differences in morphology and physiology. A similarly remarkable shift occurs in the amoeba *Dictyostelium discoideum* and related dictyostelids in their so-called social life cycle, leading to the formation of spores through aggregation of vegetative trophozoites and the development of multicellular fruiting bodies. Although dictyostelids undergo symbioses with various bacteria, chlamydiae have only recently been found to be associated with this host. Here we report the isolation of a *Dictyostelium giganteum* strain naturally infected with a chlamydial symbiont, identified as a novel species, *Reclusachlamydia socialis*. The symbiont is retained in all stages of the host’s social life cycle and notably lacks an extracellular form. Combining fluorescence microscopy and quantitative PCR we showed that transmission is entirely dependent on cell-to-cell contact during the host aggregation stage. The absence of an extracellular stage is further supported by transmission electron microscopy and the lack of genes essential for chlamydial developmental cycle regulation and extracellular survival. This unprecedented variation of a highly conserved developmental feature that evolved more than a billion years ago illustrates the remarkable adaptability of chlamydiae. This study adds to our understanding of endosymbiosis in the face of facultative multicellularity.

## Introduction

The phylum Chlamydiota is a group of obligate intracellular bacteria best known for major animal and human pathogens such as *Chlamydia trachomatis*. This species causes the ocular disease trachoma affecting 1.5 million humans globally ^1^ and infects the human genital tract at an estimated worldwide prevalence of 3% ^2^. The last three decades have expanded our knowledge of the Chlamydiota beyond those pathogens to the so-called environmental chlamydiae. These chlamydiae are found in diverse eukaryotic hosts and environments, natural and engineered, with molecular data suggesting the existence of hundreds of chlamydial families ^3^. The most characteristic feature of known chlamydiae is their biphasic developmental cycle. This lifestyle is best described for *C. trachomatis* and other members of the family Chlamydiaceae, and highly conserved among environmental chlamydiae ^4–6^. Briefly, an extracellular stage, the elementary body (EB), enters a host cell. Within a host-derived vacuole, termed inclusion, the EB radically changes its morphology and metabolism, and transforms into a reticulate body (RB). After multiple rounds of replication as RBs, the chlamydial progeny differentiates back to EBs and exits the host cell, via lysis or extrusion, to infect new hosts and start a new developmental cycle. Due to the challenging extracellular environment EBs possess a more rigid cell envelope and densely packed chromatin compared to their RB counterparts ^7–9^. The progression through this developmental program is well-orchestrated by a set of conserved chlamydiae-specific transcription factors ^10–12^ and metabolic pathways to meet the phase-specific needs of energy parasitism and energy conservation, respectively ^6,13^. Reconstruction of the genomic repertoire of the last common ancestor of the chlamydiae revealed that already 1-2 billion years ago, these bacteria were obligate endosymbionts equipped with the hallmark genes of an intracellular lifestyle and the chlamydial bi-phasic developmental cycle ^14^.

The strictly host-associated lifestyle has been hampering the isolation of environmental chlamydiae, and we thus lack isolates for the vast majority of chlamydial lineages ^3,15,16^. Yet, a source of chlamydial isolates in the past was the paraphyletic group of the free-living amoebae. These unicellular eukaryotes move via pseudopods and are ubiquitous in the environment ^17^. One of the most intensely studied free-living amoebae is the cellular slime mold *Dictyostelium discoideum*, affiliated to the clade Dictyostelia, i.e., dictyostelids, rooted within the Amoebozoa ^18,19^. Undoubtedly, the characteristic drawing the most attention to this microorganism is its social life cycle first described in the original report about *D. discoideum* ^20^ and known to occur in different variations in all dictyostelids studied so far ^18^. Briefly, the social life cycle involves the aggregation of tens of thousands of amoeba cells (trophozoites) upon nutrient deprivation to form a multicellular structure, the pseudoplasmodium, resembling a slug. This slug migrates towards light sources and other environmental cues and subsequently transforms into a mulitcellular fruiting body visible with the naked eye. The fruiting body is made up by a stalk lifting up spores ready to be dispersed in the environment and to found a new population ^21^. Despite its complexity, similar forms of facultative multicellularity have evolved at least seven times independently in amoeba ^22^. *D. discoideum* shows complex interaction patterns with facultative bacterial symbionts to ensure nutrient availability ^23^ and serves as a model to study host-microbe interaction with several human pathogens ^24,25^. Yet, the first molecular evidence of chlamydiae naturally infecting *D. discoideum* and other dictyostelids was only reported recently ^26^.

Here, we report on the isolation and characterization of *Dictyostelium giganteum* PALH, a close relative of *D. discoideum*, harboring a chlamydial symbiont remarkably well-adapted to the social life cycle of its amoeba host. The symbiont remains within the host cell throughout the entire dictyostelid social life cycle and depends on the host’s aggregation stage for the infection of naïve amoeba. This adaptation is also reflected in the symbiont genome by the loss of highly conserved genes involved in the canonical bi-phasic developmental cycle of chlamydiae.

## Results and Discussion

### A chlamydial symbiont stably infecting *Dictyostelium giganteum*

In an effort to isolate novel environmental chlamydiae, the amoeba isolate *D. giganteum* PALH was recovered from forest soil (Figure 1A) and identified based on its partial 18S rRNA gene sequence. Either on plates or in liquid cultures *D. giganteum* PALH entered the social life cycle. However, fruiting bodies and spores only formed on agar plates (Figure 1CD). By performing fluorescence in situ hybridization (FISH) with chlamydiae-specific probes, the presence of chlamydial symbionts in *D. giganteum* PALH trophozoites and spores was readily visible (Figure 1BE). Despite a chlamydial prevalence of 100% in *D. giganteum* PALH, the number of chlamydial cells counted per trophozoite and spore hardly exceeded 10 and 5, respectively (Figure 1BE). This stands in stark contrast to most other chlamydial species like *Parachlamydia acanthamoebae* ^27^*, Simkania negevensis* ^28^*, Rhabdochlamydia porcellionis* ^29^ or *C. trachomatis* ^13^, which are found in large inclusions harboring dozens of bacterial cells. The stable but low level of infection indicates that the chlamydial symbiont is highly infectious, and that its proliferation is tightly controlled either by the symbiont or the host.

**Figure 1:**
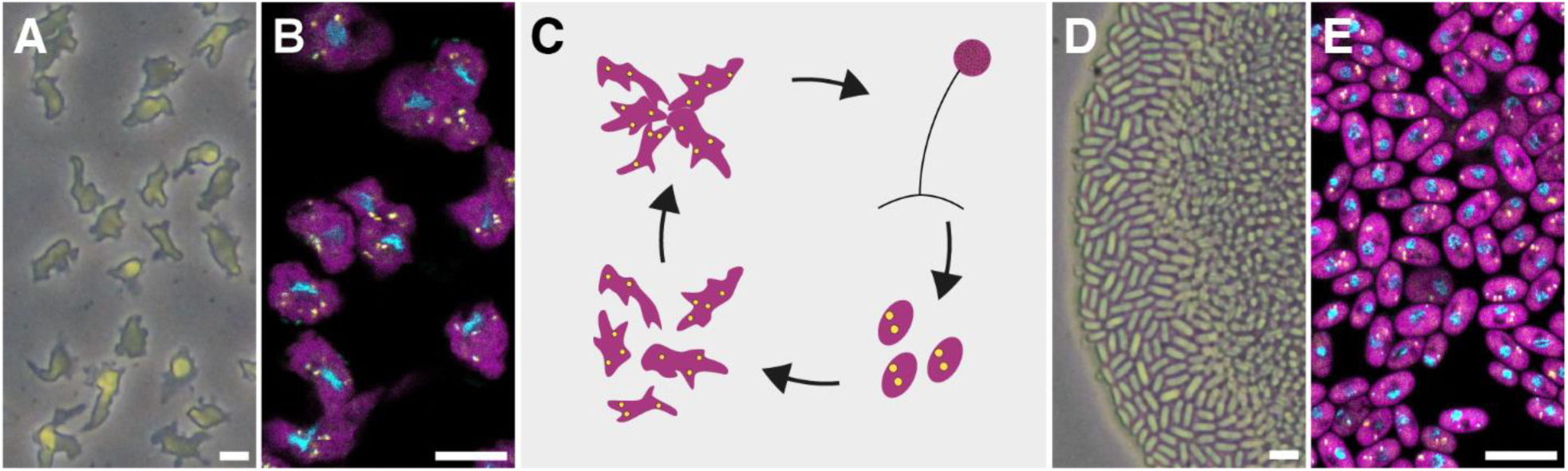
*D. giganteum* PALH naturally containing chlamydial symbionts. Light microscopy images and fluorescence in situ hybridization images of trophozoites **(A, B)** and spores **(D, E)**; nuclei are shown in cyan, amoebae in magenta, and chlamydiae in yellow. Bars, 10 µm. A scheme illustrating the presence of the symbiont during the social life cycle stages of its dictyostelid host is depicted in **(C)**; infected amoeba trophozoites and spores are shown in magenta, chlamydial symbionts in yellow.

*D. discoideum* interact with food bacteria and symbionts in manifold ways throughout their entire life cycle ^30^. In the vegetative trophozoite stage the amoeba exploit distinct pathways to discriminate between different types of food bacteria ^31^ and are able to induce endosymbiosis with food bacteria by the secretion of lectins ^32^. During multicellular stages, specialized sentinel cells are responsible for clearing pathogens via phagocytosis or extracellular traps ^33,34^. In addition, *D. discoideum* may show a farming phenotype, in which food and non-food bacteria are maintained throughout the social life cycle ^23^. This behavior is controlled by facultative intracellular *Burkholderia* spp. symbionts, which can be present in high numbers in both trophozoites and spores ^35–37^. Further, obligate intracellular *Amoebophilus* and *Procabacter* species have been detected as natural symbionts inside *D. discoideum* spores with no described fitness costs for the host ^38^. In contrast, when *D. discoideum* was artificially infected with environmental chlamydiae in the past, they either prevented spore formation or were lost during spore development (Horn et al., 2000; Michel et al., 2004). The natural chlamydial symbiont of *D. giganteum* identified here and its retention throughout the social life cycle thus point to a well-established symbiosis and a high degree of reciprocal adaptation of both partners.

### Electron microscopy reveals peculiar symbiont morphology

To elucidate the morphological features of the symbiont and its location within the amoeba host cells, we cryofixed *D. giganteum* PALH trophozoites and spores and analyzed them at the transmission electron microscope. Bacterial symbionts could be readily identified in both trophozoites and spores based on their size between 300 and 800 nm, a Gram-negative type cell wall, and the ribosomes visible in the cytoplasm (Figure 2). Unlike most chlamydiae, the symbionts were located directly in the host cytosol and not surrounded by an inclusion membrane. In other chlamydiae, this host-derived membrane is formed after host cell entry and serves to protect the bacteria from the host immune response and phagocytosis ^13^. Although wide-spread among the Chlamydiota, inclusions are not seen for all environmental chlamydiae or host cells ^3,39,40^.

**Figure 2:**
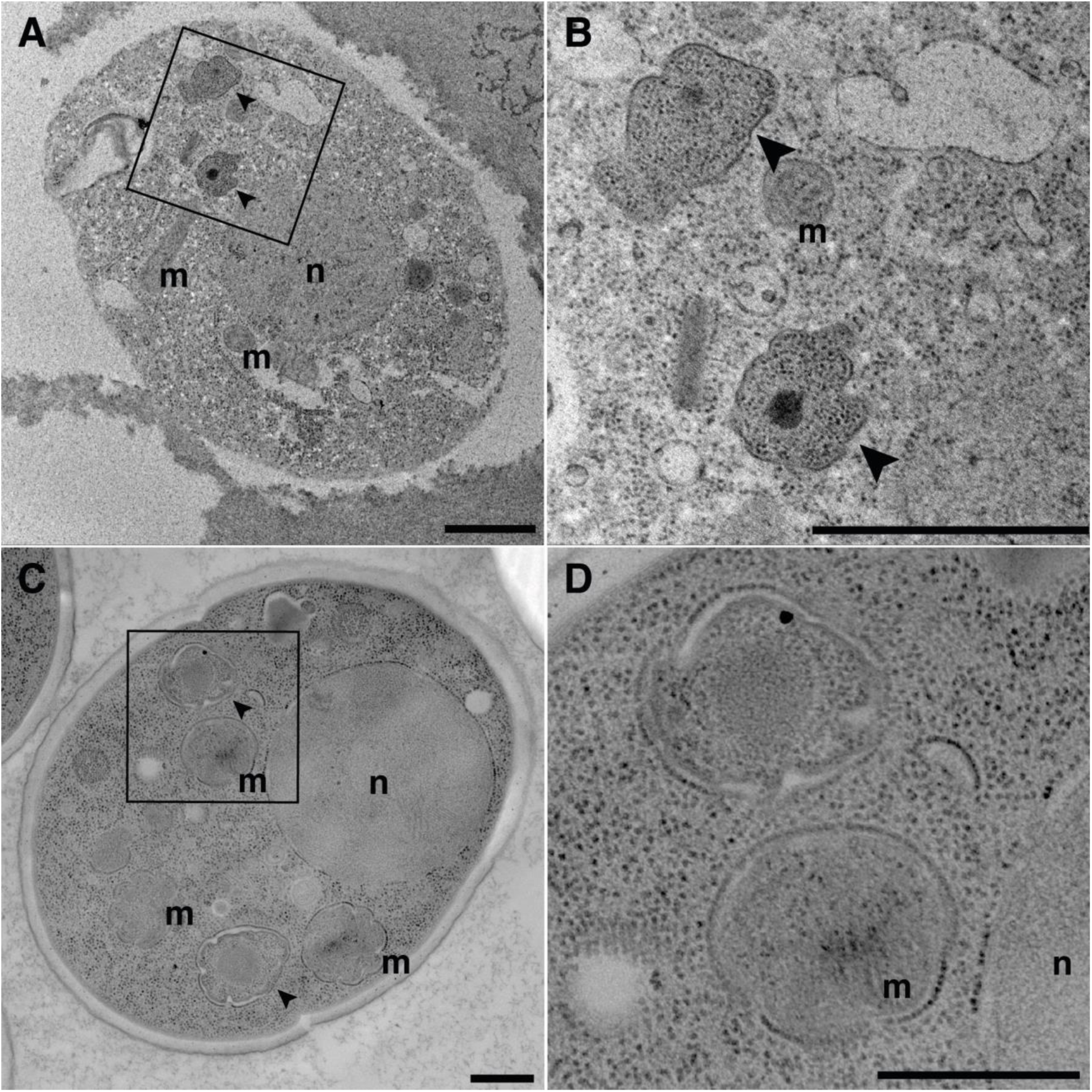
The chlamydial symbiont *Reclusachlamydia socialis* in *D. giganteum* PALH. Transmission electron microscopy images of cryofixed and freeze-substituted amoeba trophozoites and spores are shown. Overviews of a single amoebae trophozoite **(A)** and spore **(C)** harboring chlamydial symbionts (arrowheads). Rectangles indicate the area shown in detail in **(B)** and **(D)**. The chlamydial symbionts show amorphous cell shapes, contain ribosomes, a Gram-negative type cell envelope. Within the bacterial cells, nucleoid-like structures of varying electron density can be seen. Note that different staining methods were used for trophozoite and spore samples, respectively. Bars, 500 nm; n, amoeba nucleus; m, mitochondria.

The chlamydial symbionts in *D. giganteum* show a peculiar, strongly wrinkled, almost flower- or clover-like shape, in many cases containing a nucleoid-like structure of varying electron density (Figure 2). Prominent, electron-dense nucleoid-like structures are a characteristic feature of chlamydial EBs ^9,40–42^. In *C. trachomatis,* chromatin condensation is mediated by at least two histone-like proteins, Hct1 and Hct2, and is involved in the regulation of replication and transcription ^43–45^. However, in contrast to classical chlamydial EBs with their rigid and thickened cell envelope, the *D. giganteum* symbionts are of irregular shape, even more pronounced than observed for RBs of some environmental chlamydiae ^9^. This polymorphic cell shape suggests a high degree of cell envelope flexibility and a lack of rigid, stabilizing structures. The symbiont morphology is also different from described aberrant bodies, a persistent cell form induced by tryptophan depletion or peptidoglycan-targeting antibiotics in *Chlamydia* species ^46^. To our knowledge, the only chlamydial species with a comparable cell morphology was described for a chlamydial symbiont of *D. discoideum*, visualized in spores after chemical fixation ^38^.

### Symbiont genome features and phylogenetic relationship

To investigate if the observed phenotypic traits are reflected in the symbiont’s genome, we combined long and short read sequencing and obtained a closed circular genome of 1.74 Mbp with an average G+C content of 38.4%. This is well within the expected range for environmental chlamydiae ^14,16,47^. No plasmid was recovered. The genome contains one ribosomal RNA operon, 36 tRNAs and 1236 coding sequences, the latter with an average length of 1251 bp (Table S1). We next constructed a phylogenetic tree using conserved marker proteins and a data set of high-quality Chlamydiota genomes. The maximum likelihood tree placed the chlamydial symbiont in the family Rhabdochlamydiaceae, proposed previously to represent the most species-rich family within the Chlamydiota (Figure 3A, Figure S1)^48^. Members of this family were detected in arthropods such as ticks ^49^, spiders ^50,51^, and terrestrial isopods ^29^, all affiliated to the genus *Rhabdochlamydia*. Additionally, the genera *Sacchlamyda* and Renichlamydia’ were proposed based on molecular data associated with brown algae and fish, respectively ^52,53^. Average amino acid identity (AAI) values of less than 60% with other members of the Rhabdochlamydiaceae support the classification of the symbiont as a new genus within this family (Figure S1)^54,55^. We thus propose the novel name *Reclusachlamydia socialis* for the chlamydial symbiont of *D. giganteum* PALH (see Text S1 for a formal species description). In our phylogenomic analysis, members of the genus *Rhabdochlamydia* are the closest described and cultured relatives of *R. socialis*. The symbiont forms a monophyletic clade with a metagenome assembled genome (MAG) originating from wastewater, which is however only moderately related (60% AAI, Figure 3A).

**Figure 3:**
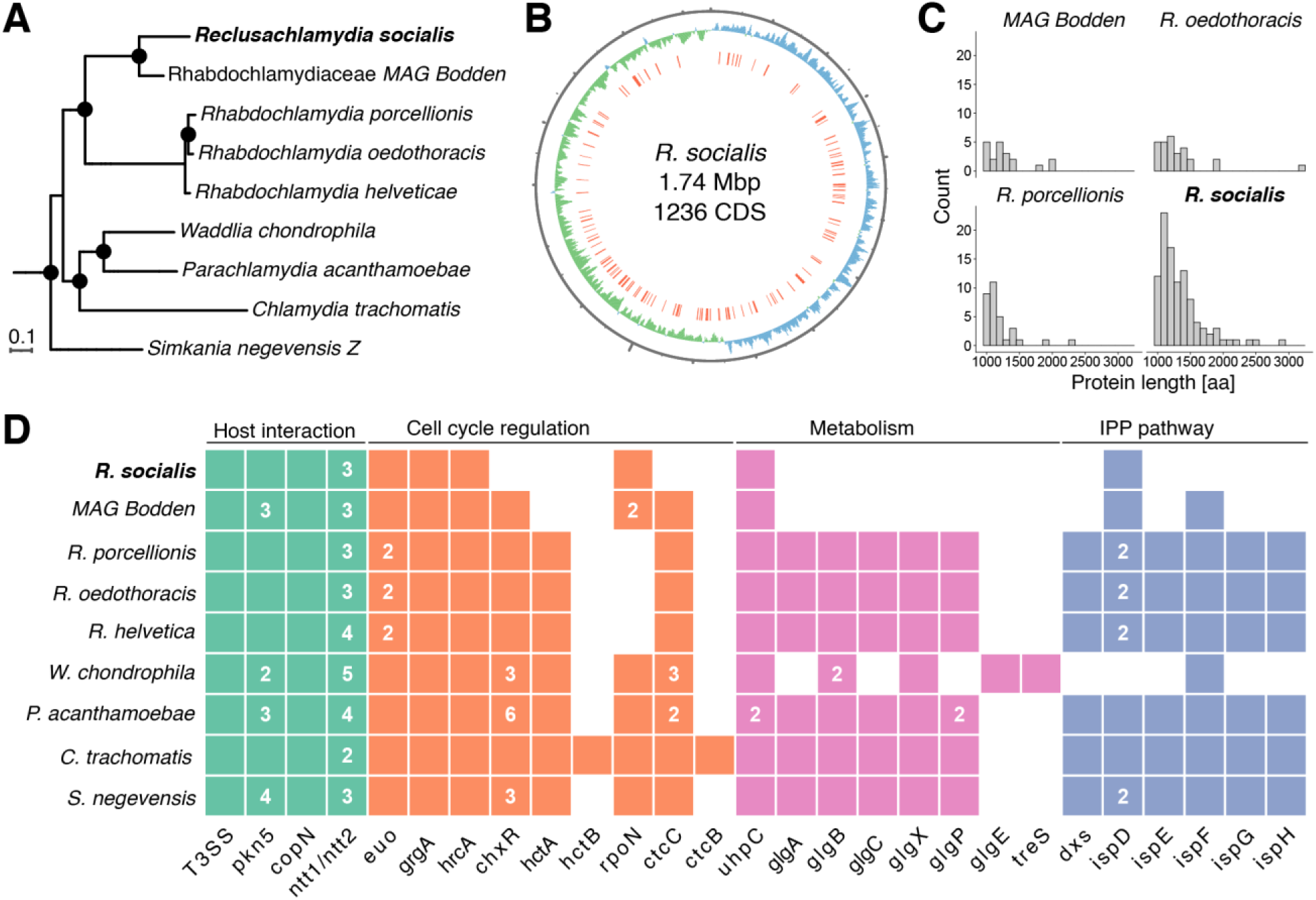
Genome features of the D. giganteum symbiont R. socialis. **(A)** Phylogenetic relationship of R. socialis *with other members of the family Rhabdochlamydiaceae based on maximum likelihood* phylogenetic *analysis of 43 conserved marker genes.* R. socialis *is the first member of the novel genus* Reclusachlamydia *and represents a sister branch of a metagenome assembled genome (MAG) from a wastewater sample. Members of the genus* Rhabdochlamydia *are the closest described and cultured relatives of* R. socialis*. The full tree is available as Data S1. **(B)** Genomic map of* R. socialis. C*oordinates* in black, *GC skew* in *green and blue; the orange bars denote the position of genes encoding large proteins (>1000 aa). **(C)** Length distribution of proteins longer than 1000 aa encoded in the genomes of selected members of the Rhabdochlamydiaceae. **(D)** Presence or absence of hallmark chlamydial genes in* R. socialis *and representative Chlamydiota species. Presence and the number of orthologs is indicated by a tile and the depicted number, respectively*.

### Large eukaryote-like proteins of *Reclusachlamydia socialis*

Upon initial inspection of the *R. socialis* genome, we noticed that 102 of the total 1236 genes (8.3%) encode unusually large proteins (> 1000 amino acids), which is unusual for small genomes and more than three times the amount reported for other members of the *Rhabdochlamydiaceae* (Figure 3BC). Of these, 91 were annotated as hypothetical proteins but we identified tetratricopeptide repeats, ankyrin repeats and L domain-like stretches in 48%, 14% and 11% of the proteins, respectively. These domains are often part of so-called eukaryote-like proteins, which are found primarily encoded in the genomes of eukaryotes and host-associated bacteria but rarely in free-living microbes. They are assumed to be secreted to the host cytosol and may interfere with host cellular processes such as cell cycle control, ubiquitination and phagocytosis ^56–58^. The importance of eukaryote-like proteins for bacteria replicating in amoebae is underlined by their high prevalence across several bacterial phyla ^57,59^.

Additionally, we identified protein prenyl transferase domains in 8 of the 102 large proteins and 15 in total. Prenyl transferases are involved in the post translational modification of proteins in eukaryotes, by specifically binding to CAAX motifs at the N-terminal end of proteins and attaching isoprenoids to the cysteine residue, which leads to the localization of the modified proteins to lipids and membranes ^60^. This CAAX motif is often found in members of the Ras superfamily of small GTPases. Small host GTPases are recruited and used for manipulation of cellular pathways by chlamydial pathogens (reviewed in ^61^). In *D. discoideum* this superfamily includes Rac proteins, such as RacG, RacJ, and RacH ^62^. RacH has been shown to influence bacterial infections in *D. discoideum* in at least two, contradicting ways. Firstly, it is a driver of the acidification of phagosomes, thus deletion of *racH* enhances intracellular growth of *Legionella pneumophila* and *Mycobacterium marinum* in *D. discoideum* ^63,64^. Conversely, RacH mediates the non-lytic transmission of *M. marinum* via ejectosomes during cell-to-cell contact ^65,66^. Eight out of the 15 putative prenyl transferases encoded in the *R. socialis* genome are predicted to be secreted by the type III secretion system (T3SS) and therefore likely function within the amoeba host cell. The manipulation of host Ras GTPases by symbiont-encoded prenyl transferases could be an efficient way of host-symbiont interaction.

### *R. socialis* lacks vital genes for extracellular survival

Similar to other obligate endosymbionts, chlamydiae experienced substantial gene loss during the millions of years of evolution as strictly host-associated bacteria. Yet, their core genome is remarkably conserved ^14,16^. Many of the core genes reflect the obligate intracellular lifestyle and are involved in the interaction with the eukaryotic host cell. The most prominent feature is the chlamydial T3SS and its affiliated effectors ^67^. The presence of all relevant structural genes and additional effectors, such as the serine/threonine protein kinase CopN and the pseudokinase Pkn5 in the genome of *R. socialis* suggest a functional type III secretion system. Similarly, the two nucleotide transporters NTT1 and NTT2, as well as the glucose-6-phosphate transporter UhpC, involved in energy parasitism, nucleotide, and glucose acquisition from the host cell ^6,13,68^ are encoded in the genome of *R. socialis* (Figure 3D, Table S1).

Of note, *R. socialis* lacks all of the known pathways for isoprenoid synthesis. The deoxyxylulose 5-phosphate (DXP) pathway is virtually absent although it is well conserved across the domain Bacteria including the Chlamydiota (Figure 3D). Additionally, none of the genes of the alternative mevalonic acid (MVA) pathway used by the chlamydial species *Waddlia chondrophila* and generally found in mammals and yeast ^69,70^ could be detected in *R. socialis*. Isoprenoids are essential cellular components involved in critical functions such as electron transport and peptidoglycan synthesis ^71^. A dependency on host-derived isoprenoid precursors has been reported for the obligate intracellular pathogen *Rickettisa parkerii* ^72^ but represents a novel type of parasitism among chlamydiae.

Progression through the bi-phasic developmental cycle is a tightly regulated process in all known members of the *Chlamydiota*. The best-known major regulator, *early upstream ORF* (EUO), is expressed during the initial infection stage, and its downregulation coincides with transformation to EBs ^73^. This process further relies on the cell cycle regulators GrgA and HrcA ^74^, all of which are encoded in the genome of *R. socialis* (Figure 3D, Table S1). Conversely, the genome of *R. socialis* lacks the genes for AtoC/AtoS, which form a two-component signal transduction system activating the σ54 transcription factor that regulates genes involved in the RB-to-EB transition ^75,76^. This coincides with the loss of ChxR, a transcriptional regulator of mid- and late cycle genes functioning in e.g., glycogen and nucleotide metabolism, modification of the inclusion membrane, and the T3SS ^10,77^.

The extracellular EB stage of chlamydiae demands protection and condensation of the DNA, which is facilitated by two histone-like proteins HctA and HctB in *C. trachomatis* ^43–45^. In contrast to *hctB*, *hctA* is conserved across most environmental chlamydiae ^16^. Yet, the genome of *R. socialis* lacks both genes (Figure 3D, Table S1). It is thus either unable to pack its chromatin or employs a yet undescribed mechanism, as suggested by transmission electron microscopy (Figure 2). To release chromatin from both HctA and HctB, *C. trachomatis* depends on the expression of IspE, a central enzyme of the DXP pathway missing in the *R. socialis* genome ^45,78^.

The extracellular chlamydial EB stage relies on efficient energy storage mediated by glycogen. Glycogen synthesis is thus considered a hallmark of chlamydiae, and all known chlamydiae encode one of two glycogen synthesis pathways ^79,80^. Yet, none of the enzymes of these pathways is encoded in the *R. socialis* genome (Figure 3D). This suggests the lack of the capability to produce and utilize this storage compound, which should severely impact the symbiont’s ability to survive outside the host cell.

We noted that the MAG Bodden seems to exhibit a very similar pattern of gene loss, suggesting that this is a common feature of this clade. Genome reduction is a process common in endosymbiosis, however, the genes lost in *R. socialis* compared to other chlamydiae indicate a reduced role or complete loss of the extracellular phase of the canonical chlamydial developmental cycle.

### Transmission of *R. socialis* is independent of extracellular chlamydiae

The absence of several hallmark genes involved in the biphasic chlamydial developmental cycle, particularly in the formation of the extracellular EB stage, led us to investigate how *R. socialis* spreads and establishes new infections. To this end, we used rifampicin to generate an aposymbiotic *D. giganteum* host culture. We subsequently tested whether it was possible to reinfect symbiont-free amoeba using supernatant from infected amoeba cultures or lysed symbiont-containing amoeba cells. Despite following well-established protocols, reinfection of aposymbiotic amoeba failed repeatedly (data not shown). We thus resorted to survey and compare infection dynamics of *R. socialis* in aposymbiotic, symbiotic, and mixed (1:5; symbiotic:aposymbiotic) *D. giganteum* populations. We monitored amoeba and symbiont total abundance in both culture supernatant and cellular fraction, as well as *R. socialis* prevalence.

*D. giganteum* cell numbers increased by a factor of ∼230 in all populations during the course of the experiment and were not significantly different between aposymbiotic, symbiotic, and mixed populations at any of the time points analysed (Figure 4A, Table S2). Nearly identical growth in the presence and absence of *R. socialis* indicate that the symbiont does not impose considerable fitness costs on its amoeba host under the monoxenic culture conditions used. This was also reflected in the maximum growth rate of all populations, which was not significantly affected by the presence of the symbiont (Figure S2). Consistent with host cell growth, the number of chlamydial genome copies measured by digital PCR increased continuously in the cellular fraction of symbiotic and mixed populations over the course of the experiment and reached similar levels in both after 72 h incubation (Figure 4B). Of note, the number of chlamydial genome copies detected at this time point in the supernatant of symbiotic and mixed populations did not differ from the aposymbiotic population (non-parametric ANOVA, p= 0.871; Figure 4B). This indicates the absence of extracellular chlamydiae in the culture medium. In parallel, we used FISH to monitor the prevalence of chlamydiae in the mixed amoeba populations. We observed a significant increase from 29.9% (SD=4.8%) infected amoeba cells at 48 h to 39.5% (SD=4.5%) at 72 h (t-test; t(15.9)=-4.35, p = 0.000498; Figure 4CD).

**Figure 4:**
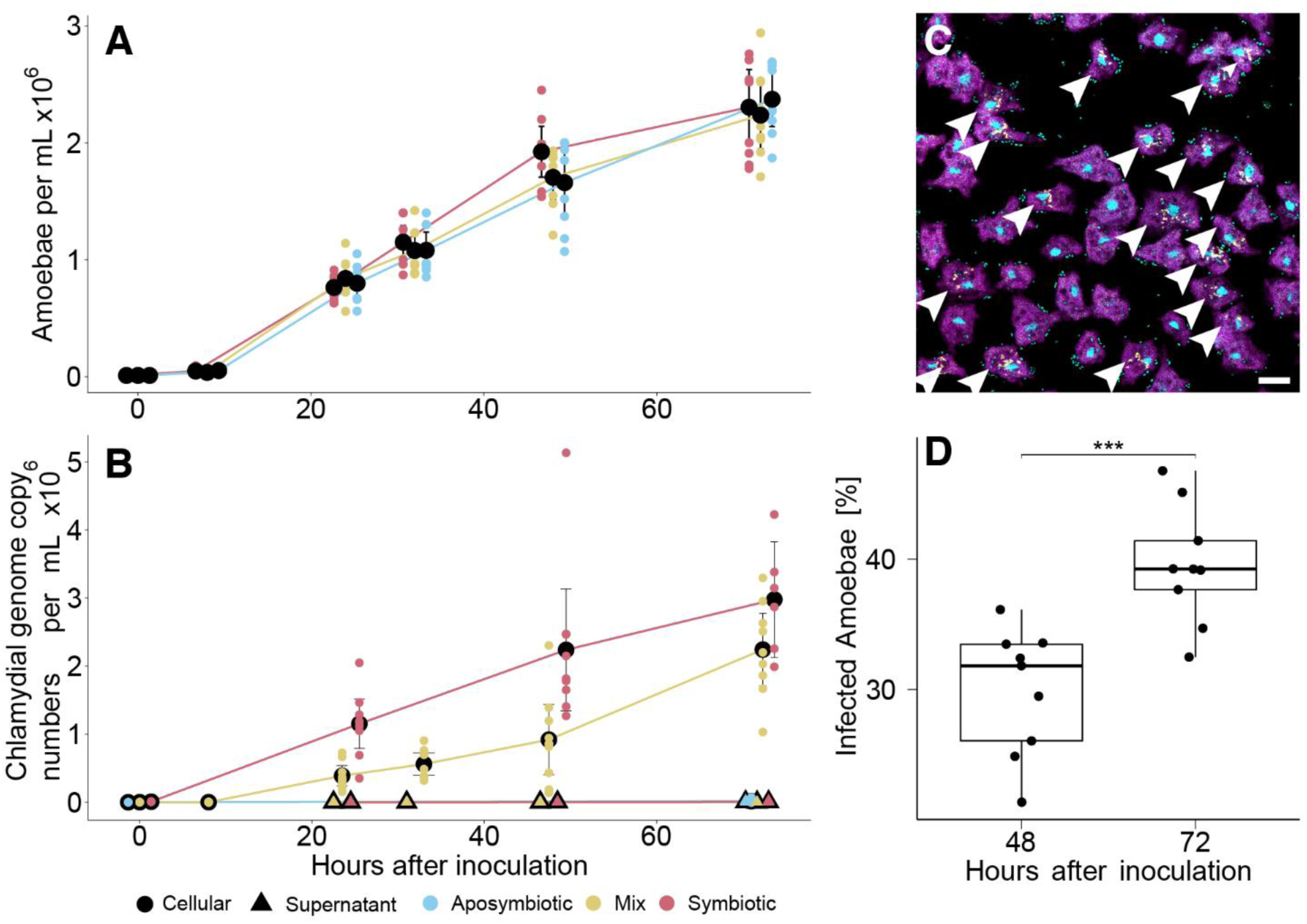
Transmission of *R. socialis* is independent of extracellular chlamydiae. Symbiotic (red), aposymbiotic (blue) or mixed (yellow; 1:5, symbiotic:aposymbiotic) *D. giganteum* PALH populations were allowed to grow in the presence of an excess of *K. aerogenes* as a food source for 72 hours. Amoeba cell numbers were monitored using a cell counter **(A)**. Chlamydial genome copy numbers were determined by dPCR in cellular fractions (circles) and supernatant (triangles), respectively **(B)**. Mean values or replicates are shown in black or color, respectively. Amoeba growth rate was not significantly affected by the presence of *R. socialis* (P > 0.05; Kruskal-Wallis test; Figure S2). Chlamydial prevalence in mixed populations was quantified 48 and 72 hours after inoculation by FISH **(C)** and increased significantly during this period (***, P < 0.001, t-test) **(D)**. Each dot represents a replicate. Boxes and error bars depict the interquartile range and the 95% confidence interval, respectively. In the fluorescence images, amoebae are depicted in magenta, chlamydiae in yellow, nuclei and food bacteria *K. aerogenes* in cyan. Infected amoebae are marked with white arrowheads. Bar, 10 µm.

Taken together, the increase of *R. socialis* cell numbers and prevalence in the mixed populations in the absence of detectable extracellular chlamydiae suggests that the infection of new host cells is independent of an extracellular stage of *R. socialis*. This would leave cell-to-cell transmission as the only option for *R. socialis* to infect naive amoeba host cells. This infection route would demand intimate contact of individual amoeba cells, a behavior frequently observed in dictyostelids, especially during the initiation of the multicellular stages ^18^.

### Infection depends on cell-to-cell contact between host cells

Exploiting the direct contact between host cells for transmission is a behavior known from human and amoeba pathogens, such as *Listeria* and *Mycobacteria* species ^66,81^. To investigate the importance of cell-to-cell contact for the transmission of *R. socialis* we co-incubated symbiotic and aposymbiotic *D. giganteum* populations under conditions leading to rapid amoeba aggregation and determined chlamydial prevalence after a short incubation period (Figure 5AC). To distinguish between originally symbiotic and aposymbiotic populations, amoeba were stained differentially using fluorescent live stains prior to co-incubation. A second set of cultures was set up identically, except that the populations were separated by a cell culture insert including a filter membrane. This filter is permeable for chlamydiae but efficiently inhibits migration of amoebae and thus hampers cell-to-cell contact between symbiotic and aposymbiotic amoeba (Figure 5BD). The size-selective retention of amoeba by cell culture inserts has been used for the isolation of chlamydiae in the past ^27^. Hence, the cell culture inserts were no barrier for EBs of the chlamydial symbiont *P. acanthamoebae* infecting *Acanthamoeba terricola* (formerly *A. castellanii* Neff) used as positive control for the extracellular transmission route in our experiment (Figure S3). At the end of the incubation period, we removed the cell culture insert containing the symbiotic amoeba. We then sampled the originally aposymbiotic population and examined the prevalence of *R. socialis* using FISH (Figure 5CD). Originally aposymbiotic *D. giganteum* cells that were allowed to freely interact with their symbiotic counterparts showed a chlamydial prevalence of 35.4% (SD=5.6%), which can be explained by the transmission of *R. socialis* between these populations. Conversely, chlamydial prevalence in originally aposymbiotic amoeba physically separated from the symbiotic population was with 3.5% extremely low (SD=1.8%; t-test; t(9.7)= 16.2, p < 0.0000001; Figure 5E). In the absence of detectable extracellular EBs, this low percentage of newly infected amoeba most likely resulted from direct interactions with the low number of symbiotic amoeba, which still passed the filter (1.7% of all cells). We thus concluded that transmission of *R. socialis* is dependent on physical contact between its host cells.

**Figure 5:**
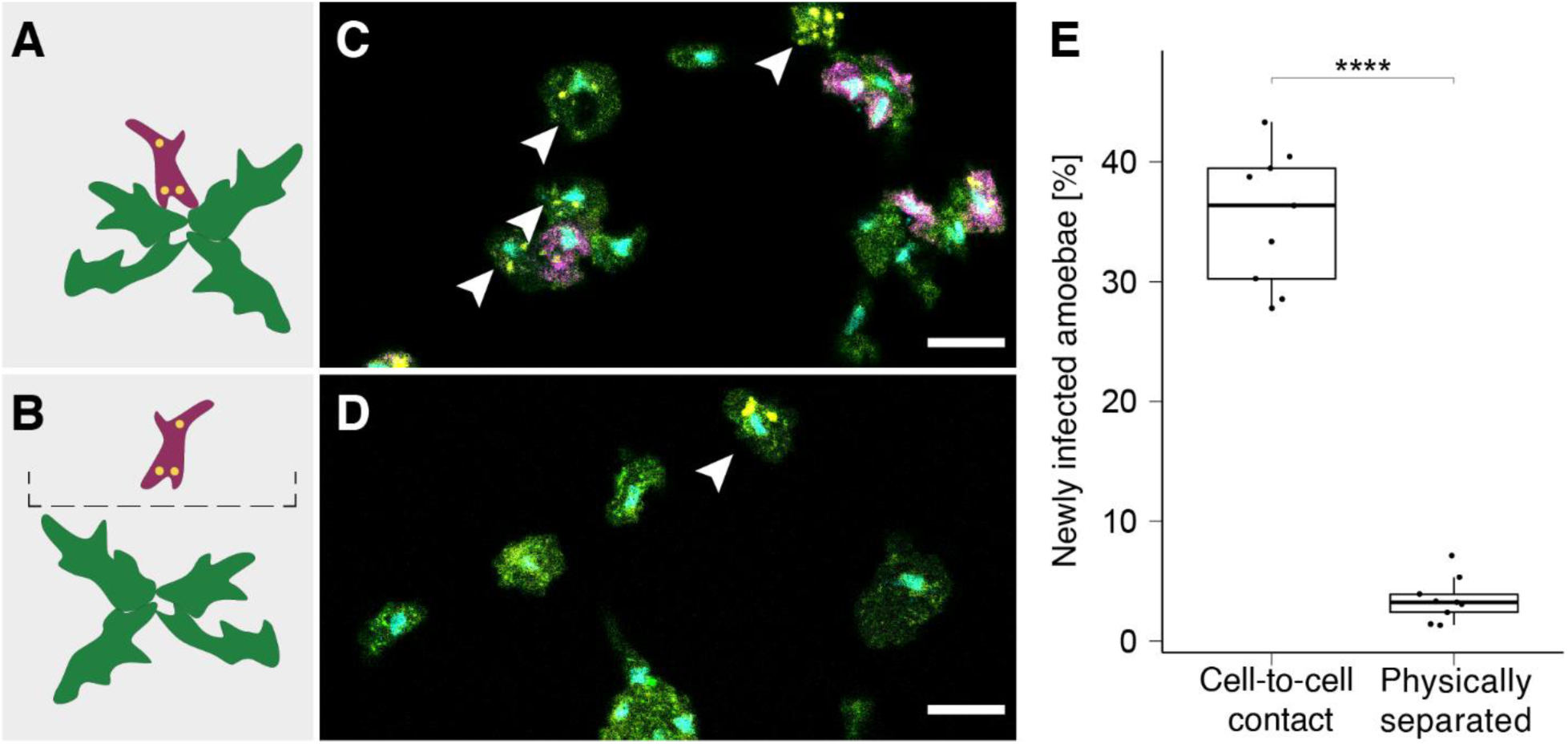
Transmission of *R. socialis* requires cell-to-cell contact between amoeba host cells: *D. giganteum* PALH populations were differentially stained with CellTracker dyes and incubated for 48h, followed by FISH to visualize *R. socialis*. Originally aposymbiotic amoeba (green) and symbiotic amoeba (magenta, chlamydiae in yellow) could either interact freely **(A, C)** or were separated physically by a filter membrane permeable for bacteria **(B, D)**. At the end of the incubation period, the number of amoeba newly infected with *R. socialis* was determined by fluorescence microscopy **(C, D)**. Non-separated populations of *D. giganteum* showed a significantly higher prevalence of chlamydiae in originally aposymbiotic cells than the physically separated populations **(E)** (****, P < 0.001, t-test). Each dot represents a replicate. Boxes and error bars depict the interquartile range and the 95% confidence interval, respectively. In the fluorescence images, newly infected amoeba are marked with white arrowheads. Bars, 10 µm.

For known chlamydiae, the infection of new host cells primarily depends on the uptake of extracellular EBs. Yet, if uptake is blocked experimentally, direct cell-to-cell transmission has been reported for *C. trachomatis* through tunneling nanotubes in human cell lines ^82^. Similar to tunneling nanotubes, *D. discoideum* exhibits temporary cell fusion, particularly during aggregation, called anastomosis ^83^. These structures facilitate the exchange of cytoplasm and even mitochondria between neighboring cells, albeit at a low rate (Bloomfield et al., 2019). *R. socialis* hijacking *Dictyostelium* anastomosis during the social life cycle could thus represent a route for cell-to-cell transmission. However, the mechanism behind this process remains to be elucidated.

## Conclusion

The chlamydiae represent one of the evolutionary oldest groups of strictly host-associated bacteria. Their ability to infect eukaryotic cells and their intracellular lifestyle can be traced back to a last common ancestor that lived around 1 billion years ago (Dharamshi et al., 2023). Here we show that one of the most conserved features of all known chlamydiae, their biphasic developmental cycle including an infective extracellular stage, shows remarkable variations in a novel chlamydial species, *R. socialis*, that infects the social amoeba *D. giganteum*.

We present evidence that the chlamydial symbiont *R. socialis* does not have an extracellular EB stage: The symbiont lacks (i) morphologically clearly distinct developmental forms, (ii) genes for extracellular survival and developmental cycle regulation, and (iii) detectable extracellular cells. The symbiont is retained through the dictyostelid social life cycle in an infected amoeba population and is dependent on cell-to-cell contact between host cells for the infection of naïve amoeba.

This represents a remarkable adaptation of a chlamydial symbiont to its host and raises the question of evolutionary causes and consequences. It is tempting to speculate that the long-term co-existence of *R. socialis* with a unicellular host frequently undergoing multicellular stages might have relaxed the dependency on an extracellular chlamydial form. This would in turn have favored an alternative transmission route in which the symbionts eventually rely on aggregation of their host cells. While the mechanism facilitating direct cell-to-cell transmission is still unclear, *R. socialis* notably encodes putative effector proteins with the potential to manipulate host cellular processes involved in the cell-to-cell transmission of *M. marinum* in *D. discoideum* (Gerstenmaier et al., 2015; Hagedorn et al., 2009).

The social life cycle of dictyostelids poses a barrier for maladapted intracellular bacteria ^33,34,84^. *R. socialis*, however, is well able to persist during the social life cycle and the spore stage. This type of vertical transmission over extended evolutionary time periods is known to lead to reduced virulence of chlamydiae (Herrera et al., 2020). More generally, vertical transmission increases cooperation between symbiotic partners (Bull et al., 1991), often resulting in obligate symbioses in which one or both partners strongly depend on the other (Husnik & Keeling, 2019). Here, we did not observe an effect of *R. socialis* on amoeba fitness with respect to host growth rate, yet host fitness is strongly context dependent (Noh et al., 2018). Bacterial symbionts of microbial eukaryotes can function in nutritional or protective symbiosis ^85–88^. In dictyostelids, this is exemplified by the symbioses between *D. discoideum* and *Paraburkholderia* species, which decrease host growth in vegetative cells but increase the probability of spore survival after germination ^23,89–92^.

The micobiome of dictyostelids in the wild is diverse and includes among other bacteria various chlamydial lineages ^26,38^. Studying these symbionts will help to further understand microbe-host interactions during and evolutionary consequences of facultative host multicellularity.

## Acknowledgements

We thank Jeffrey Blanchard for providing the soil sample for amoeba isolation, Siegfried Reipert for assistance with electron microscopy, Lisa Dobinger for help with the preparation of samples for fluorescence microscopy, Alejandro Manzano-Marin for help with genome assembly, Flo Panhölzl for help with R, and Tamara Haselkorn, Susanne DiSalvo, and Thierry Soldati for their helpful advice. Genome sequencing was performed at the Vienna Biocenter Core Facilities and the Joint Microbiome Facility of the University of Vienna and the Medical University of Vienna. The Life Science Compute Cluster LiSC at the University of Vienna provided the high-performance computing infrastructure for this study. M.H. acknowledges support from the Austrian Science Fund (FWF) [DOI 10.55776/DOC69] and the FWF Cluster of Excellence COE7 [DOI 10.55776/COE7]. A.C. was supported by the Austrian Science Fund FWF (DOI 10.55776/P32112).

## Author contributions

Conceptualization, Methodology: L.H., P.A., M.H.; Investigation, L.H., P.A., N.C.; Resources, Data Curation, A.C.; Writing – Original Draft, L.H.; Writing – Review & Editing, all authors. Funding Acquisition, M.H., A.C.

## Supplemental information

Document S1. Text S1, Figures S1–S3, Table S2

Data S1. Phylogenetic tree in Newick tree format calculated with genomes listed in Table S3.

Table S1. Excel file containing comprehensive information on the annotation of *R. socialis*.

Table S3. Excel file containing information on the genomes used for calculating the phylogenetic tree.

Table S4. Excel file containing information on the genomes used for calculating orthogroups.

## Material & Methods

### Isolation and maintenance of amoeba and bacteria

*D. giganteum* PALH was isolated from soil samples from Harvard Forest, Massachusetts, USA. An aliquot was placed on non-nutrient agar (NNA) plates (1 g/L sodium citrate, 0.4 mM CaCl_2_, 4 mM MgSO_4_, 2.5 mM KH_2_PO_4_, 15 g/L agar) and overlayed with a suspension of live *E. coli*. After incubation at 20°C and the onset of amoebae growth, trophozoites were picked from the NNA plate and transferred to a fresh *E. coli*-covered NNA plate. Eukaryotic primers TAReuk454FWD (5′-CCAGCASCYGCGGTAATTCC-3′) and TAReukREV3 (5′-ACTTTCGTTCTTGATYRA-3′) were used to amplify and sequence an approximately 520 bp region of the eukaryotic 18S rRNA gene^93^.

*D. giganteum* PALH was kept in monoxenic liquid cultures in Page’s amoeba saline buffer (PAS; 1 g/L sodium citrate, 0.4 mM CaCl_2_, 4 mM MgSO_4_, 2.5 mM KH_2_PO_4_, in MiliQ water), supplemented with *E. coli* or *K. aerogenes* as food bacteria in 25 cm² cell culture flasks at 22°C. Bacteria were inoculated in 100 mL LB-Medium (10 g/L tryptone, 5 g/L yeast-extract, 10 g/L NaCl) and incubated overnight at 37°C and 180 rpm. Subsequently, the bacterial cultures were centrifuged at 5,000 x g for 5 min, resuspended in PAS buffer to a final concentration of 2.5×10^10^ cells per mL and stored at 4°C.

### Generation of aposymbiotic amoeba

NNA plates were overlayed with 200 µL rifampicin (c=2.5 mg/mL) and 200 µL *E. coli* (2.5×10^10^ cells/mL). A single *D. giganteum* PALH fruiting body was picked with a sterile pipette tip and placed in the middle of a freshly prepared plate, which was allowed to dry and then incubated at 20°C until new fruiting bodies were formed. Fruiting bodies picked from this plate were used to repeat the procedure twice. The loss of the symbiont was confirmed by chlamydiae-specific FISH and PCR. Aposymbiotic *D. giganteum* PALH were subsequently propagated using the conditions described above.

### Transmission electron microscopy

*D. giganteum* PALH trophozoites infected with *R. socialis* were grown in 25 cm² flasks in PAS buffer supplemented with *E. coli*. At the phase of nascent aggregation, the trophozoites were harvested by centrifugation at 1500 x g for 3 min. Spores infected with *R. socialis* were grown on SM/5 plates, collected with PAS buffer in the plate lids and centrifuged at 1500 x g for 3 min. Trophozoite and spore pellets were transferred into high-pressure freezing type A carriers (3 mm in diameter, 200 μm in depth) and covered with 20% bovine serum albumin (BSA) and type B carriers (Leica Microsystems, Austria). Before use, the inner surfaces of the carriers were coated with 1-hexadecene. The mounted sample carrier sandwiches were then frozen at approximately 2000 bar using the HPM100 high-pressure freezer (Leica Microsystems, Austria). Freeze substitution was performed in an automated freeze substitution system AFS2 (Leica Microsystems, Austria). Carriers containing high-pressure frozen samples were placed on 1 mL liquid nitrogen frozen 1% OsO_4_ in acetone in 2 mL cryotubes. The following temperature program was used for freeze substitution under agitation: 2 h from -105 to -95 °C, 40 h at -95°C, 2 h from -95 °C to -90 °C, 5 h at -90 °C, 2 h from -90 °C to -60 °C, 2 h at -60 °C, 4 h from -60 °C to 20 °C. Samples were infiltrated with mixtures of 100% acetone and epoxy resin Agar 100 (Agar Scientific Ltd., Stansted, UK) according to the following schedule: 3:1 for 1 h, 1:1 for 1.5 h, 1:3 overnight. The samples were then transferred into silicon embedding molds and infiltrated with low-viscosity resin for 4 h. The resin was polymerized in an oven at 63 °C for 28 h. Ultrathin sections of 70−90 nm thickness were prepared with a Leica ultramicrotome (EM UC7, Leica Microsystems, Austria) using a diamond knife (Diatome, Nidau, Switzerland) and mounted on 200 mesh copper grids. The grids holding the ultrathin sections of trophozoite or spore pellets were mounted on a grid holder and single drop stained with gadolinium (III) acetate for 25 min (trophozoites) or 4% neodymium (III) acetate hydrate for 50 min (spores). Subsequently, the samples were rinsed and immersed in water, single drop post-stained with 3% lead citrate for 8 min (in a covered petri dish with NAOH pellets to scavenge lead carbonate) and then washed again with water. The stained ultrathin sections were examined with a Zeiss Libra 120 transmission electron microscope.

### Fluorescence *in situ* hybridization

Aliquots of *D. giganteum* cell suspensions were placed on teflon-coated microscope slides and incubated for 15 minutes at room temperature to allow attachment of the trophozoites. The PAS buffer droplet was then replaced with 20 µL of 4% (w/v) paraformaldehyde (PFA) in PAS buffer and incubated for 10 min at room temperature. Fixation was terminated by removing PFA/PAS and briefly washing each well with 50 µL water. The fixed cells were covered with 10 µL of hybridization buffer (900 mM NaCl, 20 mM Tris-HCl, 0.01% SDS, 25 % formamide) mixed with 1 µL of the respective probes and hybridized at 46°C for 2 h in an atmosphere saturated with hybridization buffer. The slides were then incubated in washing buffer (20 mM Tris-HCl, 0.25 mM EDTA and 0.149 M NaCl) at 48°C for 15 min and washed in ice-cold water for 10 s. The probes used in this study were Chla-232 (5’-TAG CTG ATA TCA CAT AGA -3’), Chls-523 (5’-CCTCCGTATTACCGCAGC -3’) ^94^ each 5’-labelled with the fluorescent dye Cy3 for the detection of the chlamydial symbiont, and Euk-516 (5’-GGA GGG CAA GTC TGG T -3’) 5’-labelled with Cy5 for the detection of amoeba ^95^. To stain DNA, all wells were covered with 10 µL 4′,6-diamidino-2-phenylindole (DAPI) in deionized water (c = 1 µg / mL) for 3 min and subsequently washed for 7 min in 99% ethanol. Citifluor AF1 (Electron Microscopy Sciences, USA) was used to mount a coverslip. Slides were analyzed using a Leica TCS SP8 X confocal laser scanning microscope equipped with a 93x glycerol objective and the Leica application suite X software (v3.7.6).

### Growth and infection experiments

To assess the maximum growth rates of symbiotic, aposymbiotic, and mixed (1:5, symbiotic:aposymbiotic) *D. giganteum* PALH populations, trophozoites in log phase were incubated at a starting cell concentration of 10^4^ cells per mL in PAS buffer supplemented with *K. aerogenes* in 25 cm^2^ cell culture flasks (Fisher Scientific Austria GmbH, Austria) at 22°C. At defined timepoints, amoeba cells were detached from the bottom of the cell culture flasks using cell scrapers. Aliquots were taken and frozen for subsequent DNA extraction (250 µL) and measurement of amoeba concentrations (10 µL) using the LUNA-FX7™ Automated Cell Counter (Logos Biosystems, South Korea). For microscopy, 40 µL aliquots were applied to microscope slides and allowed to attach prior to chemical fixation and FISH with chlamydiae- and eukaryote-specific probes. To analyse the supernatant for extracellular chlamydiae, 1 mL aliquots of the supernatant were taken, homogenized using 25G syringes to break possible bacterial cell aggregates, and filtered through 1.2 µm filters to remove amoebae. Samples were frozen for subsequent DNA extraction and digital PCR.

To study the effect of physical contact between amoeba host cells on symbiont transmission, log phase symbiotic and aposymbiotic trophozoites were washed to remove food bacteria and stained with CellTracker deep red and green, respectively, according to the manufacturer’s protocol (Thermo Fisher Scientific Inc., USA). Trophozoites were then inoculated at a concentration of 5×10^5^ cells/mL and a ratio of 1:5 of symbiotic to aposymbiotic amoeba either mixed in 12-well plates or separated using Nunc™ Cell Culture Inserts in Carrier Plate Systems with a pore size of 3 µm (Thermo Fisher Scientific Inc., USA). After 48 h of incubation at 22 °C, amoeba were harvested from the wells and transferred to a microscope slide. Chlamydial prevalence was assessed by FISH followed by DAPI staining. Positive controls consisting of *Acanthamoeba terricola (*formerly *A. castellanii* Neff) infected with *P. acanthamoebae* were also assessed but the amoeba were derived from cultures grown in TSY medium (30 g/L trypticase soy broth, 30 g/L yeast extract). Infection with *P. acanthamoebae* was performed 48 hours before the start of the co-incubation, at an MOI of 200. Chlamydiae and amoebae were mixed, centrifuged at 150 x g for 15 min, and the supernatant was exchanged after 3 h to remove excess chlamydiae.

### Symbiont quantification by digital PCR

Aliquots of frozen amoeba cells were thawed, centrifuged at 10,000 x g for 10 min and resuspended in 200 µL PAS buffer. DNA was then extracted using the QIAamp 96 Virus QIAcube HT Kit (QIAGEN, NL) according to the manufacturer’s protocol. The efficiency of the extraction was tested using the Qubit system (Thermo Fisher Scientific Inc., USA). Digital PCR was performed using the QX200 Droplet Digital PCR System (Biorad, USA). The final PCR reaction mixture contained 1x QX200™ ddPCR™ EvaGreen Supermix, SigF2 (5’-CRGCGTGGATGAGGCAT-3’) and SigR2 (5’-TCAGTCCCARTGTTGGC-3’) primers (0.18 pM) ^96^, 2 µL of the respective DNA sample and ddH_2_O in a total volume of 22 µL. Droplet generation was performed according to the manufacturer’s protocol. PCR was performed with the following steps: enzyme activation at 95°C for 5 min, 40 steps of denaturation at 95°C for 30 s and annealing at 59°C for 1 min, signal stabilization at 4°C for 5 min, and a final step at 90°C for 5 min. The droplet suspension was subsequently analyzed using the QX200 droplet reader combined with the QX Manager standard edition software (v 1.2). To distinguish positive from negative droplets, a fluorescence amplitude threshold of 6000 was set uniformly for all samples.

### Genome sequencing and assembly

Symbiotic *D. giganteum* PALH cultures were incubated in PAS buffer supplemented with *E. coli*, until the bottom of twelve 175 cm² cell culture flasks was confluent. The supernatant was replaced with fresh PAS buffer without food bacteria, and the amoebae were detached using a cell scraper. The detached amoebae were transferred to 50 mL tubes and centrifuged at 5000 x g for 10 minutes. The resulting pellet was resuspended in PAS buffer, transferred to 2 mL Eppendorf tubes prefilled with 0.5 mL glass beads (d=0.25-0.5mm) and vortexed horizontally at 2700 x g for 2 min. The disrupted amoeba cells were further homogenized using a 25G needle and passed sequentially through 5 µm and 1.2 µm pore size filters to remove host cell debris and nuclei. The Wizard® HMW DNA Extraction Kit (Promega, USA) was then used to extract DNA for long read sequencing according to the manufacturer’s protocol. Library preparation was performed at the Vienna Bio Center Core Facilities using the Multiplex Ligation Sequencing Kit and the Native Barcoding Kit (Oxford Nanopore Technologies, Oxford, UK). Long-read sequencing was performed on an ONT Flongle system. Libraries for short read sequencing were prepared using the NEBNext Ultra II FS DNA Library Prep Kit for Illumina, pooled equimolarly, and sequenced using an Illumina MiSeq instrument (2 x 300bp, 600 cycles).

Short reads were trimmed using cutadapt 3.5 ^97^ to avoid adapter contamination and low quality sequences. Long reads were trimmed using Porechop 0.2.4 ^98^ and corrected using Canu 2.0 <u>(Koren et al., 2017)</u>. To assemble the *R. socialis* genome, all reads were used in a hybrid approach using Unicycler 0.4.8 ^99^. The resulting single chlamydial contig was then rotated and the origin of replication determined using Ori-finder 2022 ^100^ and originx ^101^.

### Annotation, comparative genomics, and phylogenetic analysis

Gene calling and annotation was performed using bakta 1.9.1 ^102^. The coding sequences called by bakta were the basis for the subsequent comparative genome analysis. InterProScan 5 ^103^ was used for protein domain identification ^104^. The prediction of effector proteins was performed using EffectiveT3 ^105^. AAI was calcuated using the enve-omics tool box with minimum alignment length 0 aa, minimum identity 20 % and minimum score 0 bits (http://enve-omics.ce.gatech.edu/aai/) ^106^. To understand patterns of gene loss and gain, orthofinder 2.5.5 was used ^107^ to construct orthogroups with a comprehensive dataset of 174 chlamydial genomes and high-quality MAGs (Table S4). Synteny regions, regions coding for long proteins, and the GC skew were visualized using circos ^108^.

For phylogenomics, CheckM ^109^ was used to produce a concatenated alignment of 43 marker proteins including the homologs of 171 chlamydial and 89 Planctomycetota-Verrucomicrobiota outgroup representatives (Table S3). This alignment was passed to IQ-TREE 2.2.6 to compute a maximum likelihood tree ^110^. Modelfinder plus ^111^ selected LG+C20+R4 as optimal model, and the final tree was calculated with ultrafast bootstraps ^112^, and the (Shimodaira-Hasegawa) SH-like approximate likelihood ratio test ^113^ with 1000 repeats each.

### Statistical Analysis

All statistical analyses were performed using RStudio Statistical Software (v2024.04.2+764; Posit team, 2024)

### Data availability

The *R. socialis* genome sequence and annotation is available under Bioproject no. PRJEB79376, accession no. ERZ24898538.

