## Supplemental File 1 for "The adaptation of chlamydiae to facultative host multicellularity"

**Text S1: Description of Reclusachlamydia socialis, gen. nov., sp. nov.**

Based on the complete genome sequence (accession no. ERS21338603) and the intracellular visualization by fluorescence microscopy and electron microscopy, we refer to the chlamydial symbiont described in this paper as ‘Reclusachlamydia socialis’ [Re.clu.sa.chla.my.di.a, Lat. fem. n., reclūsa, hermit; N.L. fem. n. Chlamydia, taxonomic name of a bacterial genus; socialis. Lat. gen. n. socialis, referring to the characteristic behavior of the host organism, a dictyostelid]. ‘Reclusachlamydia socialis’ PALH is a strictly intracellular symbiont within the cytoplasm of the social amoeba *Dictyostelium giganteum*, isolated from soil samples of Harvard Forrest (Massachusetts, USA). The bacteria are non-motile, only found within and not cultivable outside their amoeba host cells, and able to persist during the amoeba social life cycle. The morphology of ‘Reclusachlamydia socialis’ in amoeba trophozoites and spores resembles the morphology of unclassified chlamydial symbionts described in *Dictyostelium discoideum* [1]. The symbiont cells appear wrinkled and folded, occasionally with an area of distinct electron density in the centre of the cell. The cells range in size from 300 to 800 nm. Based on phylogenetic analysis of chlamydial core genes and the 16S rRNA gene sequence, ‘Reclusachlamydia socialis’ forms a distinct lineage within the family Rhabdochlamydiaceae, order Chlamydiales, phylum Chlamydiota. The currently closest cultured representative is *Rhabdochlamydia porcellionis* (50.08% AAI, 89.98% 16S rRNA sequence identity).

| clean1924 | OFTM358 | 1159587-4 | Bin65_10 | 1021942-13 | Ga0074140 | NC_groundwater_600_49_25 | L024912 | PMV_8.15 | co.spades.CONCOCT.1kb_142 | 3300020970-2 | 3300005529-103 | nzgw621 | Gw_Bodden_bin_68 | RSOCI | 1095360-24 | palsa_1444 | SZUA-160 | 1096102-30 | KR126_chlam_1 | 3300009709-46 | ZH-Jul18-154 | time.spades.CONCOCT.1kb_023 | LacPavin_0818_WC40_MAG_42_9 | LacPavin_0419_WC53_MAG_40_24 | LacPavin_0419_WC53_MAG_41_8 | Amazon_FNV_2010_10_1_1_2 | RhabHel | ROED_W744 | RHAB15C | |
|---|---|---|---|---|---|---|---|---|---|---|---|---|---|---|---|---|---|---|---|---|---|---|---|---|---|---|---|---|---|---|
| 100 | 49 | 51 | 51 | 51 | 52 | 52 | 50 | 50 | 51 | 51 | 51 | 51 | 50 | 48 | 50 | 51 | 50 | 49 | 50 | 50 | 49 | 49 | 50 | 50 | 50 | 49 | 50 | 50 | 49 | clean1924 |
| 49 | 100 | 51 | 52 | 50 | 52 | 52 | 50 | 51 | 51 | 51 | 51 | 51 | 50 | 48 | 50 | 50 | 51 | 51 | 51 | 49 | 49 | 49 | 50 | 51 | 51 | 50 | 50 | 50 | 50 | OFTM358 |
| 51 | 51 | 100 | 57 | 54 | 57 | 56 | 53 | 53 | 56 | 54 | 55 | 55 | 58 | 54 | 60 | 60 | 54 | 53 | 53 | 52 | 51 | 52 | 53 | 54 | 54 | 52 | 53 | 53 | 53 | 1159587-4 |
| 51 | 52 | 57 | 100 | 54 | 57 | 57 | 53 | 54 | 56 | 55 | 56 | 55 | 54 | 52 | 55 | 55 | 53 | 53 | 54 | 52 | 52 | 53 | 54 | 54 | 54 | 53 | 53 | 54 | 54 | Bin65_10 |
| 51 | 50 | 54 | 54 | 100 | 54 | 54 | 52 | 52 | 54 | 53 | 53 | 53 | 52 | 50 | 52 | 53 | 51 | 51 | 51 | 51 | 51 | 51 | 52 | 53 | 53 | 51 | 51 | 52 | 52 | 1021942-13 |
| 52 | 52 | 57 | 57 | 54 | 100 | 64 | 55 | 55 | 57 | 56 | 57 | 56 | 54 | 52 | 55 | 55 | 54 | 53 | 54 | 53 | 52 | 52 | 55 | 55 | 55 | 53 | 54 | 54 | 54 | Ga0074140 |
| 52 | 52 | 56 | 57 | 54 | 64 | 100 | 54 | 55 | 57 | 56 | 56 | 55 | 54 | 51 | 55 | 55 | 53 | 53 | 54 | 53 | 53 | 52 | 54 | 55 | 55 | 53 | 54 | 54 | 54 | NC_groundwater_600_49_25 |
| 50 | 50 | 53 | 53 | 52 | 55 | 54 | 100 | 52 | 53 | 53 | 53 | 53 | 52 | 50 | 52 | 52 | 53 | 52 | 52 | 51 | 50 | 50 | 52 | 52 | 52 | 51 | 51 | 51 | 51 | L024912 |
| 50 | 51 | 53 | 54 | 52 | 55 | 55 | 52 | 100 | 54 | 53 | 53 | 53 | 52 | 50 | 52 | 52 | 55 | 54 | 55 | 51 | 50 | 51 | 52 | 53 | 53 | 52 | 52 | 53 | 52 | PMV_8.15 |
| 51 | 51 | 56 | 56 | 54 | 57 | 57 | 53 | 54 | 100 | 59 | 60 | 59 | 53 | 51 | 55 | 54 | 53 | 53 | 53 | 53 | 53 | 53 | 55 | 55 | 56 | 56 | 55 | 56 | 56 | co.spades.CONCOCT.1kb_142 |
| 51 | 51 | 54 | 55 | 53 | 56 | 56 | 53 | 53 | 59 | 100 | 59 | 57 | 53 | 50 | 54 | 53 | 52 | 52 | 52 | 53 | 52 | 52 | 53 | 54 | 55 | 55 | 55 | 56 | 56 | 3300020970-2 |
| 51 | 51 | 55 | 56 | 53 | 57 | 56 | 53 | 53 | 60 | 59 | 100 | 63 | 53 | 51 | 54 | 54 | 53 | 52 | 53 | 53 | 52 | 53 | 54 | 55 | 55 | 57 | 57 | 57 | 57 | 3300005529-103 |
| 51 | 51 | 55 | 55 | 53 | 56 | 55 | 53 | 53 | 59 | 57 | 63 | 100 | 53 | 51 | 54 | 54 | 53 | 53 | 53 | 53 | 52 | 53 | 54 | 54 | 55 | 55 | 56 | 56 | 55 | nzgw621 |
| 50 | 50 | 58 | 54 | 52 | 54 | 54 | 52 | 52 | 53 | 53 | 53 | 53 | 100 | 60 | 61 | 61 | 51 | 51 | 52 | 50 | 50 | 50 | 52 | 52 | 52 | 51 | 51 | 52 | 51 | Gw_Bodden_bin_68 |
| 48 | 48 | 54 | 52 | 50 | 52 | 51 | 50 | 50 | 51 | 50 | 51 | 51 | 60 | 100 | 56 | 56 | 49 | 49 | 49 | 49 | 49 | 50 | 50 | 50 | 50 | 50 | 50 | 50 | 50 | RSOCI |
| 50 | 50 | 60 | 55 | 52 | 55 | 55 | 52 | 52 | 55 | 54 | 54 | 54 | 61 | 56 | 100 | 73 | 52 | 51 | 52 | 51 | 50 | 51 | 52 | 52 | 53 | 52 | 52 | 52 | 52 | 1095360-24 |
| 51 | 50 | 60 | 55 | 53 | 55 | 55 | 52 | 52 | 54 | 53 | 54 | 54 | 61 | 56 | 73 | 100 | 52 | 51 | 52 | 51 | 51 | 51 | 52 | 53 | 53 | 51 | 51 | 52 | 52 | palsa_1444 |
| 50 | 51 | 54 | 53 | 51 | 54 | 53 | 53 | 55 | 53 | 52 | 53 | 53 | 51 | 49 | 52 | 52 | 100 | 70 | 63 | 51 | 50 | 50 | 52 | 52 | 52 | 51 | 51 | 51 | 51 | SZUA-160 |
| 49 | 51 | 53 | 53 | 51 | 53 | 53 | 52 | 54 | 53 | 52 | 52 | 53 | 51 | 49 | 51 | 51 | 70 | 100 | 62 | 50 | 50 | 50 | 51 | 51 | 52 | 50 | 51 | 51 | 51 | 1096102-30 |
| 50 | 51 | 53 | 54 | 51 | 54 | 54 | 52 | 55 | 53 | 52 | 53 | 53 | 52 | 49 | 52 | 52 | 63 | 62 | 100 | 51 | 50 | 50 | 52 | 52 | 52 | 51 | 52 | 52 | 51 | KR126_chlam_1 |
| 50 | 49 | 52 | 52 | 51 | 53 | 53 | 51 | 51 | 53 | 53 | 53 | 53 | 50 | 49 | 51 | 51 | 51 | 50 | 51 | 100 | 67 | 58 | 54 | 54 | 55 | 51 | 51 | 52 | 51 | 3300009709-46 |
| 49 | 49 | 51 | 52 | 51 | 52 | 53 | 50 | 50 | 53 | 52 | 52 | 52 | 50 | 49 | 50 | 51 | 50 | 50 | 50 | 67 | 100 | 58 | 54 | 54 | 54 | 51 | 51 | 51 | 51 | ZH-Jul18-154 |
| 49 | 49 | 52 | 53 | 51 | 52 | 52 | 50 | 51 | 53 | 52 | 53 | 53 | 50 | 50 | 51 | 51 | 50 | 50 | 50 | 58 | 58 | 100 | 54 | 54 | 54 | 50 | 51 | 52 | 51 | time.spades.CONCOCT.1kb_023 |
| 50 | 50 | 53 | 54 | 52 | 55 | 54 | 52 | 52 | 55 | 53 | 54 | 54 | 52 | 50 | 52 | 52 | 52 | 51 | 52 | 54 | 54 | 54 | 100 | 75 | 65 | 52 | 53 | 53 | 53 | LacPavin_0818_WC40_MAG_42_9 |
| 50 | 51 | 54 | 54 | 53 | 55 | 55 | 52 | 53 | 55 | 54 | 55 | 54 | 52 | 50 | 52 | 53 | 52 | 51 | 52 | 54 | 54 | 54 | 75 | 100 | 65 | 53 | 53 | 53 | 53 | LacPavin_0419_WC53_MAG_40_24 |
| 50 | 51 | 54 | 54 | 53 | 55 | 55 | 52 | 53 | 56 | 55 | 55 | 55 | 52 | 50 | 53 | 53 | 52 | 52 | 52 | 55 | 54 | 54 | 65 | 65 | 100 | 52 | 53 | 53 | 53 | LacPavin_0419_WC53_MAG_41_8 |
| 49 | 50 | 52 | 53 | 51 | 53 | 53 | 51 | 52 | 56 | 55 | 57 | 55 | 51 | 50 | 52 | 51 | 51 | 50 | 51 | 51 | 51 | 50 | 52 | 53 | 52 | 100 | 87 | 86 | 87 | Amazon_FNV_2010_10_1_1_2 |
| 50 | 50 | 53 | 53 | 51 | 54 | 54 | 51 | 52 | 55 | 55 | 57 | 56 | 51 | 50 | 52 | 51 | 51 | 51 | 52 | 51 | 51 | 51 | 53 | 53 | 53 | 87 | 100 | 88 | 87 | RhabHel |
| 50 | 50 | 53 | 54 | 52 | 54 | 54 | 51 | 53 | 56 | 56 | 57 | 56 | 52 | 50 | 52 | 52 | 51 | 51 | 52 | 52 | 51 | 52 | 53 | 53 | 53 | 86 | 88 | 100 | 89 | ROED_W744 |
| 49 | 50 | 53 | 54 | 52 | 54 | 54 | 51 | 52 | 56 | 56 | 57 | 55 | 51 | 50 | 52 | 52 | 51 | 51 | 51 | 51 | 51 | 51 | 53 | 53 | 53 | 87 | 87 | 89 | 100 | RHAB15C |

**Figure S1: AAI values of Rhabdochlamydiaceae proteomes, related to Figure 3.** AAI was calculated using the AAI calculator on http://enve-omics.ce.gatech.edu/aai/ [2]. Alignment options: Minimum length: 0aa, Minimum identity: 20%, Minimum score 0 bits, Minimum alignments 50. Each row and column represent a species, the value at the crosspoints show the respective AAI.

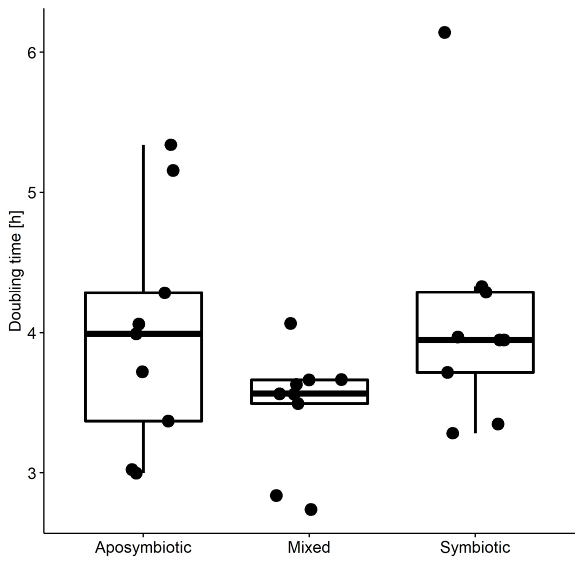


**Figure S2: Fastest doubling time of trophozoites during growth curve experiment, related to Figure 4.** *D. giganteum* PALH cultures were incubated at $10^4$ cells/ml and allowed to grow under excess of food bacteria. The fastest doubling time of trophozoites of all cultures occurred between 8 and 24 hours after inoculation. The average time for one successful cell division was 4.00 (SD=0.84), 4.11 (SD=0.84) and 3.47 (SD=0.42) hours, for aposymbiotic, symbiotic and mixed cultures, respectively. A non-parametric ANOVA test was performed and yielded no significant difference (Chi square = 4.10, p = 0.129, df = 2) between these cultures. Each dot represents one replicate. The mean, the interquartile range and the 95% confidence interval are depicted by the horizontal line, the box and the whiskers, respectively.

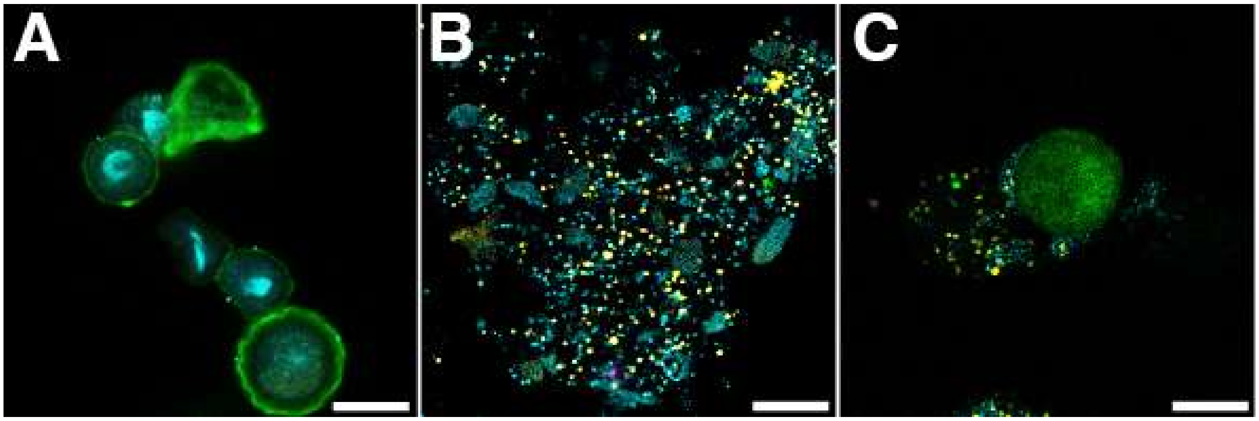


**Figure S3: Transmission of *Parachlamydia acanthamoebae* through cell-culture inserts with 3 µm pore size, related to Figure 5.** *Acanthamoeba terricola (*formerly *A. castellanii* Neff*)* trophozoites were grown in axenic medium either uninfected **(A)** or infected **(B)** with *Parachlamydia acanthamoeba* UV7. The two cultures were co-incubated but separated with a cell-culture insert of 3 µm pore size **(C)**. At the end of the experiment the presence of chlamydiae was evaluated by FISH. Formerly aposymbiotic cells are depicted in green, symbiotic cells in magenta, chlamydiae in yellow, nuclei and bacteria in cyan. Bar, 10µm.

**Table S2: Statistical comparison of quantification derived from incubations of symbiotic, aposymbiotic and mixed populations of *D. giganteum* PALH, related to Figure 4.** Trophozoites or chlamydial genome copies were quantified over the course of the incubation. Trophozoites were measured using an automated cell counter, chlamydial genome copy numbers were measured using ddPCR and chlamydiae-specific primers. Chlamydial genome copy numbers are either originating from cellular fractions of trophozoites or the supernatant. Symbiotic and mixed populations were compared by t-test and ANOVA,

| Organism | Origin | Time [h] | Variable | Compared Pop | p | p.adj | p.format | p.signif | method |
|---|---|---|---|---|---|---|---|---|---|
| Amoebae | Cellular | 8 | Cells per mL | All | 0.2349 | 0.9400 | 0.2300 | ns | Anova |
| Amoebae | Cellular | 24 | Cells per mL | All | 0.5382 | 1.0000 | 0.5400 | ns | Anova |
| Amoebae | Cellular | 32 | Cells per mL | All | 0.6725 | 1.0000 | 0.6700 | ns | Anova |
| Amoebae | Cellular | 48 | Cells per mL | All | 0.1694 | 0.8500 | 0.1700 | ns | Anova |
| Amoebae | Cellular | 72 | Cells per mL | All | 0.7141 | 1.0000 | 0.7100 | ns | Kruskal-Wallis |
| Chlamydiae | Cellular | 72 | Genomes per mL | All | 0.0002 | 0.0002 | 0.0002 | *** | Kruskal-Wallis |
| Chlamydiae | Cellular | 0 | Genomes per mL | Symbiotic - Mix | 0.0226 | 0.0450 | 0.0230 | * | T-test |
| Chlamydiae | Cellular | 24 | Genomes per mL | Symbiotic - Mix | 0.0010 | 0.0040 | 0.0010 | ** | T-test |
| Chlamydiae | Cellular | 48 | Genomes per mL | Symbiotic - Mix | 0.0019 | 0.0056 | 0.0019 | ** | Wilcoxon |
| Chlamydiae | Cellular | 72 | Genomes per mL | Symbiotic - Mix | 0.1135 | 0.2000 | 0.1135 | ns | Wilcoxon |
| Chlamydiae | Supernatant | 72 | Genomes per mL | All | 0.8711 | 0.8700 | 0.8700 | ns | Kruskal-Wallis |
| Chlamydiae | Supernatant | 24 | Genomes per mL | Symbiotic - Mix | 0.0279 | 0.0840 | 0.0280 | * | T-test |
| Chlamydiae | Supernatant | 48 | Genomes per mL | Symbiotic - Mix | 0.0244 | 0.0490 | 0.0240 | * | Wilcoxon |
| Chlamydiae | Supernatant | 72 | Genomes per mL | Symbiotic - Mix | 0.1694 | 0.1700 | 0.1690 | ns | T-test |

***References***


1. Haselkorn, T.S., Jimenez, D., Bashir, U., Sallinger, E., Queller, D.C., Strassmann, J.E., and DiSalvo, S. (2021). Novel Chlamydiae and *Amoebophilus* endosymbionts are prevalent in wild isolates of the model social amoeba *Dictyostelium discoideum*. Environ. Microbiol. Rep. 13, 708–719. https://doi.org/10.1111/1758-2229.12985.

2. Rodriguez-R, L.M., and Konstantinidis, K.T. (2016). The enveomics collection: a toolbox for specialized analyses of microbial genomes and metagenomes. PeerJ Prepr. 4, e1900v1. https://doi.org/10.7287/peerj.preprints.1900v1.
